# LAPTM4B Alleviates Pulmonary Fibrosis by Enhancing NEDD4L-Mediated TGF-β Signaling Suppression

**DOI:** 10.1101/2024.08.30.610429

**Authors:** Kai Xu, Xiaoyue Pan, Hui Lian, Yaxuan Wang, Ruyan Wan, Zhongzheng Li, Xin Pan, Yajun Li, Juntang Yang, Ivan Rosas, Lan Wang, Guoying Yu

## Abstract

Idiopathic pulmonary fibrosis (IPF) is a chronic and progressive lung disease with fatal outcome and a poorly understood pathogenesis. The lysosomal protein transmembrane 4 beta (LAPTM4B), a multi-transmembrane endo-lysosomal membrane protein, has been implicated in the pathogenesis of several diseases. However, its involvement in IPF remains unexplored. This study aimed to investigate the role of LAPTM4B in lung fibrosis and elucidate its underlying mechanisms. The results showed that LAPTM4B was significantly reduced in IPF and mouse fibrotic lungs. In vivo studies showed that the deficiency of LAPTM4B exacerbated bleomycin-induced lung fibrosis, while the restoration of LAPTM4B alleviated fibrosis. Mechanistically, LAPTM4B recruits the NEDD4 like E3 ubiquitin protein ligase (NEDD4L) to endosomes, leading to increased ubiquitin-mediated proteasomal degradation of TGFRB2 and active SMAD2/3, thereby blocking the TGF-β/SMAD signaling pathway. Overall, our data provided a novel insight for a deeper understanding of the pathogenesis of IPF, supporting the therapeutic potential of restoration of LAPTM4B as a promising therapeutic approach for the treatment of pulmonary fibrosis.

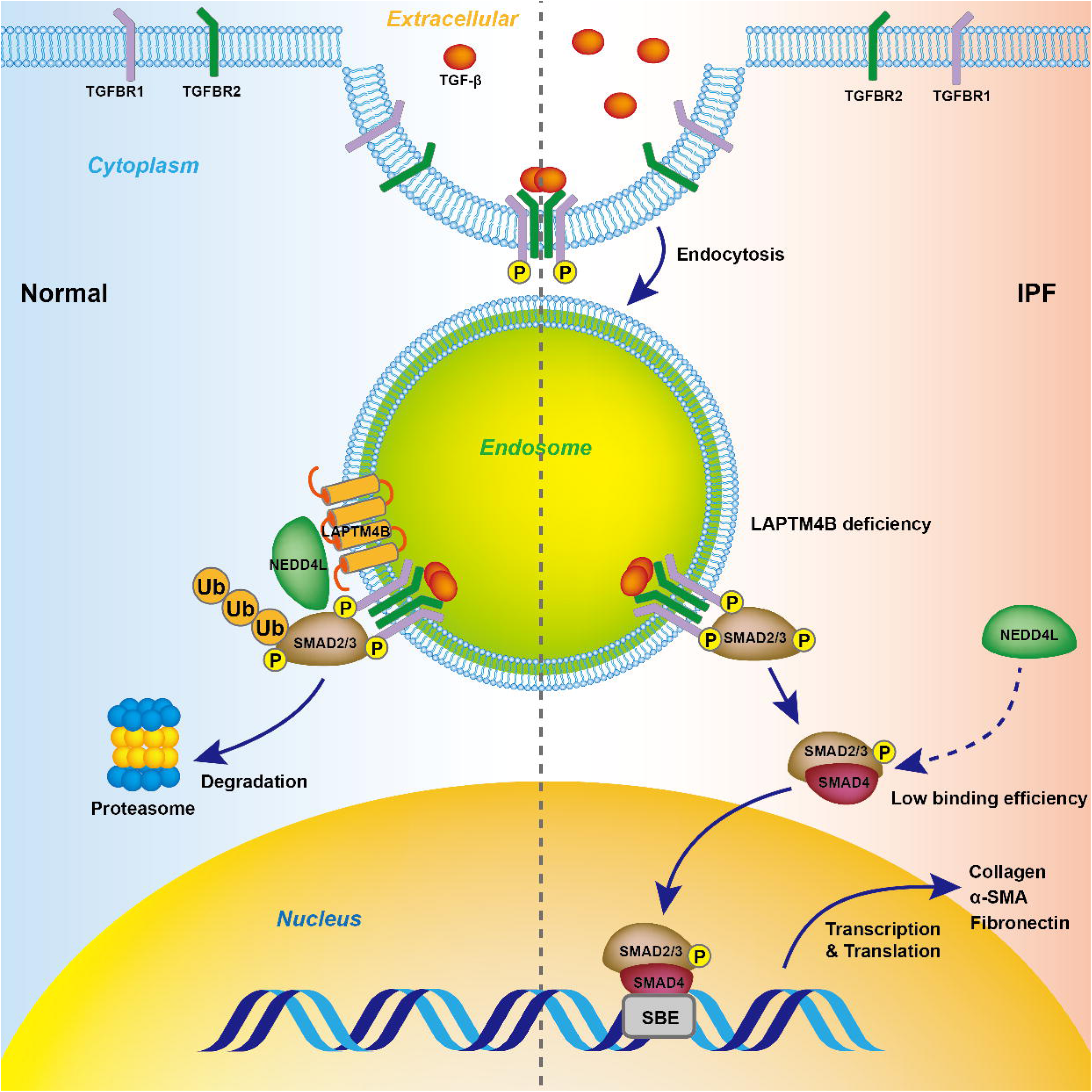

## Introduction

Idiopathic pulmonary fibrosis (IPF) is a progressive and fatal interstitial lung disease involves a complex interplay of cell types and signaling pathways (1). With a median survival time of only 3-5 years from diagnosis, IPF carries a prognosis worse than that of most malignancies (2–4). Various risk factors, including genetic predisposition, smoking, male gender predominance, and aging, have been implicated in the development of IPF (5,6). Despite extensive research efforts, the precise pathological mechanisms underlying IPF remain elusive. It is widely postulated that recurrent injuries to alveolar epithelial cells and dysregulated epithelial repair processes are pivotal, with fibroblasts serving as the primary effector cells responsible for aberrant extracellular matrix production (7,8). Currently, treatment options of IPF are limited, and management without transplant primarily revolves around supportive care measures such as oxygen therapy and pulmonary rehabilitation (9). While pharmacological therapies pirfenidone and nintedanib have been shown to mitigate disease progression, their long-term efficacy and tolerability remain uncertain (9–11). With ever-increasing incidence due to the expanding global aging population, addressing this unmet clinical challenge requires a deeper understanding of pathogenesis of IPF through continued research endeavors.

Endocytosis-mediated cellular signaling is a fundamental process that modulates cellular signal transduction via multiple molecular mechanisms (12,13). Endosomes regulate receptor internalization, recycling, and degradation, thereby modulating the transmission of extracellular signals (14,15). In addition, endosomes function as signaling platforms, recruiting and activating specific signaling molecules within endocytic vesicles, influencing intracellular signal transduction (16–18). Emerging evidence suggests that the endocytosis and trafficking of TGF-β receptors play a crucial role in regulating their fate and, consequently, the transduction of TGF-β/SMAD signaling and pro-fibrotic responses in IPF (19,20). A recent study demonstrated that Nestin, a type VI intermediate filament protein, acts as a central regulator of the intracellular vesicular trafficking system and facilitates Rab11-dependent recycling of TGFBR1, thereby sustaining TGF-β/SMAD signaling and promoting fibrogenesis in IPF (21). Despite these advances, a comprehensive understanding of endocytosis-mediated cellular signaling in IPF is essential for elucidating the pathogenesis and developing novel therapeutic strategies for the disease.

The lysosomal protein transmembrane (LAPTM) family comprises essential endo-lysosomal proteins, including LAPTM4A, LAPTM4B, and LAPTM5. Among these, LAPTM4B has been extensively studied. Previous research has demonstrated that LAPTM4B plays a critical role in regulating intracellular vesicle trafficking, sphingolipid homeostasis, nutrient sensing, and lysosomal function, thereby contributing to cellular homeostasis and signal transduction (22–25). It has been reported that LAPTM4B inhibits EGF-induced EGFR intraluminal sorting and lysosomal degradation, resulting in enhanced and prolonged EGFR signaling (26). Furthermore, LAPTM4B is upregulated in various cancers and plays a pivotal role in tumor initiation and progression, particularly in mechanisms of chemotherapy resistance (27–29). Beyond cancer research, a recent study demonstrated that LAPTM4B mitigates experimental ischemia/reperfusion injury by promoting autophagy (30). However, the involvement of LAPTM4B in fibrotic diseases has not yet been reported.

Our initial analysis suggests that LAPTM4B may play a role in the progression of pulmonary fibrosis. Building upon this hypothesis, the primary objective of this study was to investigate the involvement of LAPTM4B in the pathogenesis of IPF and to elucidate its underlying molecular mechanisms. Through this research, we aim to provide novel insights into the pathogenesis of IPF and identify potential targets for therapeutic intervention.

## Results

### LAPTM4B protein levels are reduced in IPF and bleomycin-induced fibrotic lungs

To investigate the potential role of LAPTM4B in pulmonary fibrosis, we examined LAPTM4B expression in fibrotic lungs. Histological examination revealed a notable decrease in LAPTM4B protein expression in lung sections from IPF subjects compared to non-IPF donors (Figure 1A). Consistently, immunofluorescence staining highlighted reduced LAPTM4B expression in bleomycin-treated fibrotic lungs compared to control counterparts (Figure 1B), along with an increase in the mesenchymal marker Vimentin and a decrease in the alveolar epithelial cell marker Advanced Glycation End-product Receptor (AGER). Western blot analysis confirmed the reduced level of LAPTM4B in the lung homogenates from bleomycin-challenged fibrotic lungs, accompanied by elevated Collagen1 expression (Figure 1C–D). Additionally, LAPTM4B expression was reduced in mouse acute lung injury (ALI) models induced by both bleomycin (Figure 1E&F) and lipopolysaccharide (LPS) (Figure 1G&H). However, reanalysis of the Gene Expression Omnibus (GEO) database GSE124685 revealed no significant changes in LAPTM4B mRNA levels (Supplementary Fig. 1A), and GSE150910 showed higher LAPTM4B mRNA levels in IPF lungs compared to controls (Supplementary Fig. 1B) (31,32). Similarly, Laptm4b mRNA expression was not significantly altered in bleomycin-induced fibrotic lungs (Supplementary Fig. 1C). Additionally, analyzing the single-cell RNA-seq data from the Human Lung Cell Atlas revealed that LAPTM4B is widely expressed in the lung, with relatively higher expression in airway epithelial cells, alveolar epithelial cells, and fibroblasts (Figure 1I-K) (33). This observation was further confirmed through immunofluorescence staining of normal lung tissue from *Sftpc-*tdTomato mice, where LAPTM4B was found to be distributed in puncta and partially co-localized with *Sftpc*-positive ACE2 cells (Figure 1L).

**Figure 1.**
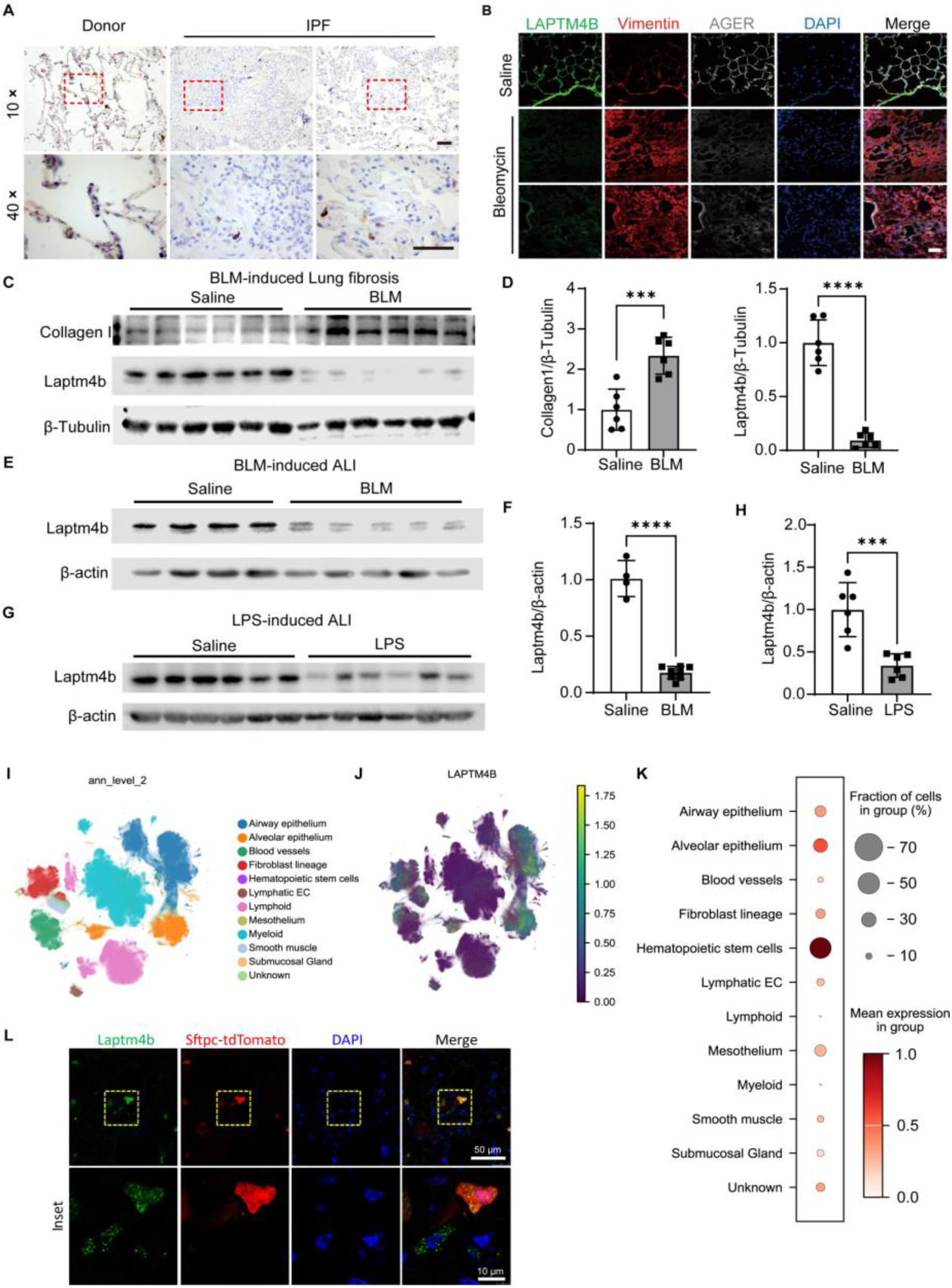
LAPTM4B levels are decreased in IPF and BLM-induced fibrotic lungs. (**A**) Representative images of IHC staining of LAPTM4B in paraffin sections of lung tissues from IPF patients and non-IPF donors. The boxed regions in images captured under 10× objective were showed in detail under 40× objective, scale bar: 50 μm. (**B**) Representative confocal images of immunofluorescent staining showing Laptm4b (green), Vimentin (red), and AGER (gray) in frozen sections of mouse lung tissues at 14 days post single-dose BLM challenge or saline treatment. Cell nuclei were counterstained with DAPI (blue). Scale bar: 50 μm. (**C**) Western blot and (**D**) densitometry analysis of Laptm4b and Collagen1 in lung homogenates from bleomycin-induced fibrotic models. β-Tubulin served as a loading control. (mean ±SD, n = 6 mice per group). (**E**) Western blot (**F**) and densitometry analysis of Laptm4b in lung homogenates from BLM-induced ALI model. β-actin served as a loading control. (mean ±SD, Saline: n = 4, and BLM: n = 5 mice). (**G**) Western blot and (**H**) densitometry analysis of Laptm4b in lung homogenates from LPS-induced ALI model. β-actin served as a loading control. (mean ±SD, n = 6 mice per group). (**I**) Uniform Manifold Approximation and Projection (UMAP) plot from the Human Lung Cell Atlas, showing the distribution of different lung cell types. Each color represents a distinct cell type. (**J**) UMAP plot and (**K**) dot plot illustrating the expression levels of LAPTM4B across various lung cell types. (**L**) Representative confocal images of immunofluorescent staining showing Laptm4b (green), and Sfptc-tdTomato (red) in mice lungs, DAPI (blue) was used to label cell nuclei. *\*\*\* p* < 0.001, *\*\*\*\* p* < 0.0001.

### AAV shRNA-mediated knockdown of Laptm4b exacerbates bleomycin-induced lung fibrosis

To investigate the impact of Laptm4b deficiency on the progression of pulmonary fibrosis, C57BL/6 mice were exposed to AAV-sh-*Laptm4b* (Figure 2A). Micro-CT scan revealed evident lung tissue lesions characterized by increased parenchymal opacity in the BLM-challenged mice, with the sh-*Laptm4b*+BLM group exhibiting even more severe lung tissue damage (Figure 2B). Consistently, histological analysis with hematoxylin-eosin (H&E) and Masson’s trichrome staining revealed no discernible alteration in lung architecture in the AAV-sh-NC+Saline and AAV-sh-*Laptm4b*+Saline groups (Figure 2C).

**Figure 2.**
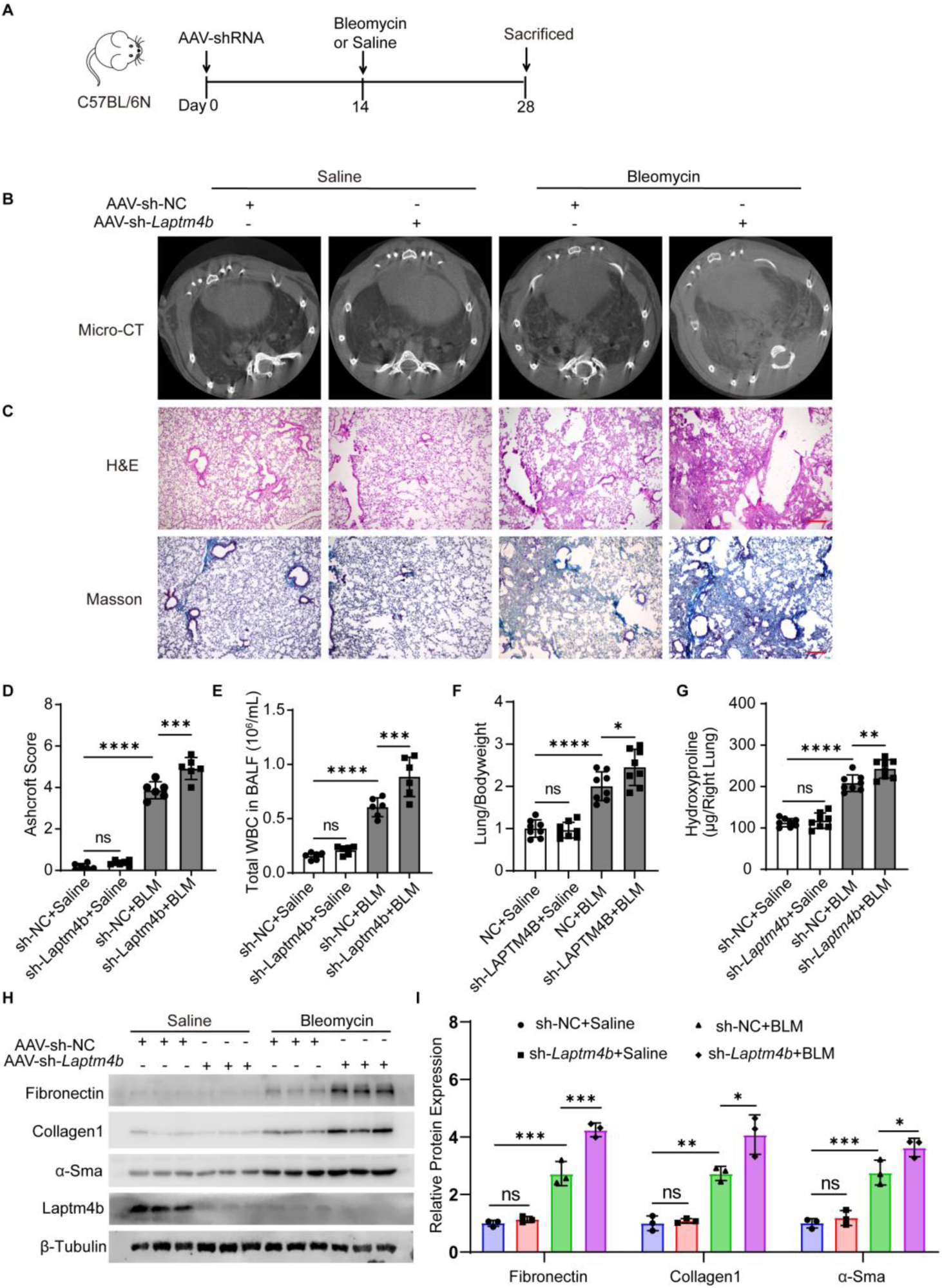
AAV shRNA-mediated Laptm4b knockdown exacerbates BLM-induced lung fibrosis. (**A**) Schematic diagram illustrating the experimental design. (**B**) Representative axial micro-CT scan images of the mouse lungs. (**C**) Representative images of hematoxylin and eosin (H&E) staining, and Masson’s trichrome staining of lung sections. Scale bar: 200 μm. (**D**) Ashcroft score analysis of H&E-stained lung sections (mean ± S.D., n = 6). (**E**) Quantification of the total number of WBC in BALF (mean ± S.D, n = 6). (**F**) Mouse whole lung weight vs body weight (mean ± S.D., n = 8). (**G**) Measurement of hydroxyproline content in the right lungs (mean ±S.D., n = 6-8). (**H**) Western blot analysis and (**I**) quantification of Fibronectin, Collagen 1, α-Sma and Laptm4b levels in mouse lung homogenates (mean ± S.D., n = 3). β-Tubulin was used as an internal control. All statistical analysis was performed using ANOVA followed by Tukey post hoc corrections. \**p* < 0.05, \*\**p* < 0.01,\*\*\**p* < 0.001, and \*\*\*\**p* < 0.0001; ns, not significant.

However, AAV-sh-*Laptm4b* exposed mice displayed more severe lung injury, and increased lung collagen deposition after the BLM challenge (Figure 2D). Additionally, there is a significant increase in the Ashcroft scores (Figure 2E), total white blood cell (WBC) counts in BALF(Figure 2F), lung-to-body weight ratios, and lung hydroxyproline content of AAV-sh-*Laptm4b* exposed mice following bleomycin challenge (Figure 2G). Western blot analysis confirmed a significant reduction in Laptm4b expression after AAV-sh-*Laptm4b* exposure and demonstrated a notable elevation in the expression of extracellular matrix (ECM) proteins Fibronectin and Collagen 1, as well as the myofibroblast marker α-SMA in the lung homogenates of AAV-sh-*Laptm4b* exposed mice following bleomycin challenge (Figure 2H&I). Collectively, these data indicate that the loss of Laptm4b exacerbates lung injury and fibrosis induced by BLM.

### LAPTM4B plays a crucial role in maintaining epithelial and fibroblast cell homeostasis by restraining TGF-β1-induced pro-fibrotic phenotypes

To assess the impact of LAPTM4B deficiency in vitro, we constructed stable LAPTM4B-knockdown A549 cell lines using two different short hairpin RNAs (shRNAs), sh-*LAPTM4B* or sh-NC, with lentiviral vectors.. Immunofluorescence analysis revealed significant changes in cell morphology in sh-*LAPTM4B* cells, accompanied by a loss of the epithelial cell marker E-cadherin and increased fluorescence intensity of Vimentin (Figure 3A). Additionally, cell migration was promoted in sh-*LAPTM4B* A549 cells, as evaluated by wound healing assay (Figure 3B&C). Western blot results further confirmed the reduced LAPTM4B expression and loss of E-cadherin, as well as increased expression of the pro-fibrotic markers N-cadherin and Vimentin in sh-*LAPTM4B* A549 cells (Figure 3D&E). Similarly, siRNA-mediated LAPTM4B knockdown induced the formation of α-SMA-positive actin fibers in human embryonic lung fibroblast MRC-5 cells (Figure 3F). Western blot analysis further confirmed the induction of Fibronectin, Collagen1, and α-SMA by siRNA-mediated LAPTM4B knockdown in MRC-5 cells (Figure 3G&H), indicating that this was not a cell type-specific result. Conversely, LAPTM4B overexpression reduced the elevation of Fibronectin, Collagen1, and α-SMA (Figure 3I&J) induced by TGF-β1. Additionally, LAPTM4B inhibited TGF-β1-induced cell contractility (Figure 3K&L), while siRNA-mediated knockdown of LAPTM4B enhanced TGF-β1-induced cell contractility in MRC-5 cells (Figure 3M&N).

**Figure 3.**
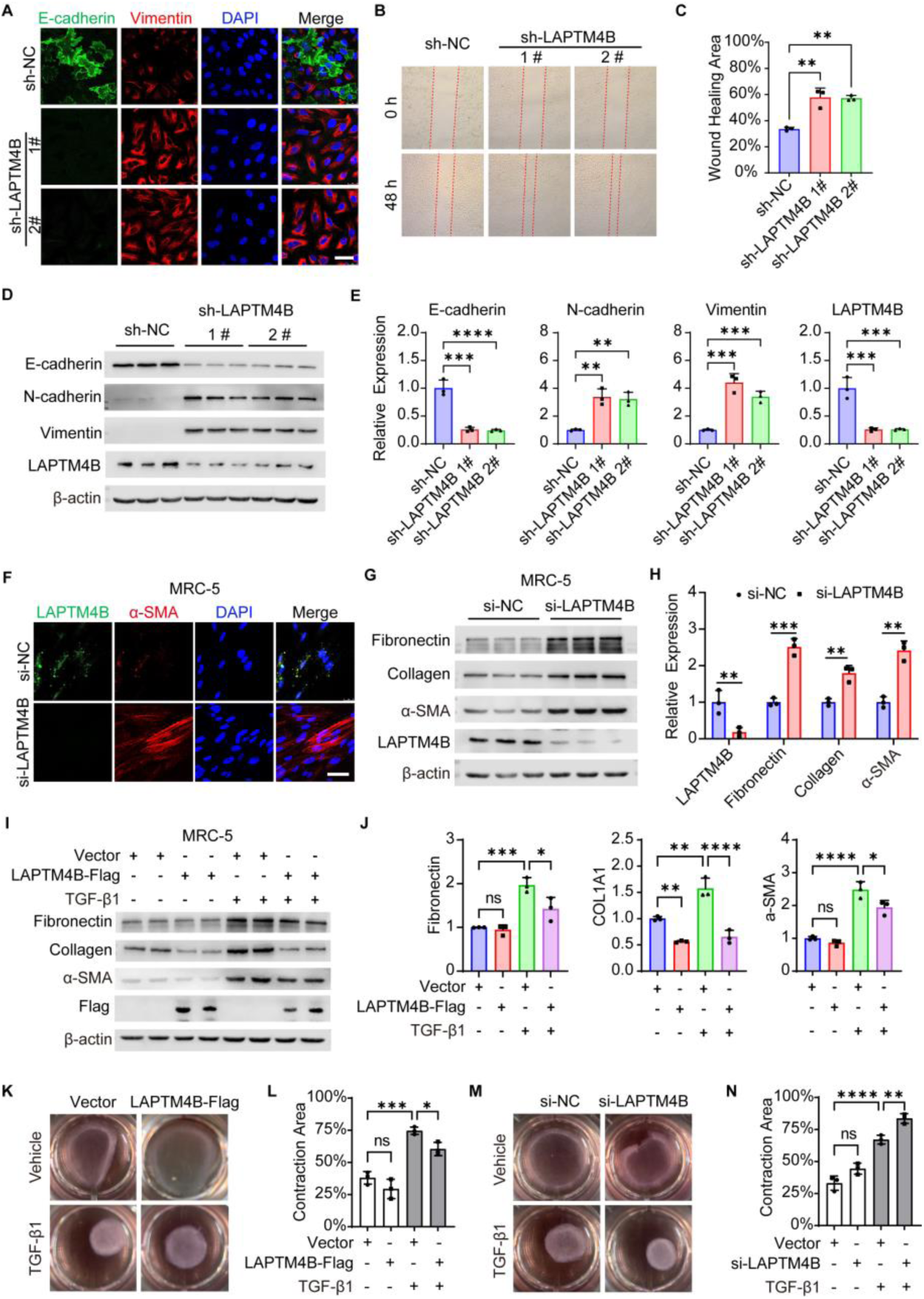
LAPTM4B plays a crucial role in maintaining the homeostasis of epithelial and fibroblast cells and attenuates TGF-β1-induced pro-fibrotic response. (**A**) Confocal images of immunofluorescent staining of E-cadherin (green) and Vimentin (red) in sh-NC and sh-MUT A549 cells. DAPI (blue) was used to label cell nuclei, scale bar: 50 μm. (**B**) Representative images of scratch assay conducted on sh-NC and sh-LAPTM4B A549 cells. The red dotted lines delineated the wound edges. (**C**) Quantification of wound closure areas from (**B)**, data presented as % of the original wound areas (mean ± S.D., n = 3). (**D**) Western blot and (**E**) densitometry analysis (mean ± SD, n = 3) of E-cadherin, N-cadherin, Vimentin, and LAPTM4B in sh-NC and sh-LAPTM4B A549 cells. β-actin was used as an internal control. (**F**) Representative confocal images of immunofluorescent staining of LAPTM4B and α-SMA in MRC-5 cells transfected with si-NC or si-LAPTM4B for 48 h. DAPI (blue) was used to label cell nuclei, scale bar: 50 μm. (**G**) Western blot and (**H**) densitometry analysis of Fibronectin, Collagen1, α-SMA, and LAPTM4B in NC and si-LAPTM4B MRC-5 cells. β-actin was used as an internal control. (**I**) Western blot and (**J**) densitometry analysis (mean ± SD, n = 3) of Fibronectin, Collagen1, and α-SMA in MRC-5 cells transfected with LAPTM4B-Flag followed by TGF-β1 treatment for 48 h. β-actin was used as an internal control. (**K**) Collagen gel contraction assay and (**L**) quantification (mean ± SD, n = 3) performed on MRC-5 cells transfected with LAPTM4B-Flag or vector control for 24 h, followed by treatment with TGF-β1 for an additional 24 h. (**M**) Collagen gel contraction assay and (**N**) quantification (mean ± SD, n = 3) performed on MRC-5 cells transfected with si-NC or si-LAPTM4B for 24 h, followed by treatment with TGF-β1 for an additional 24 h. Statistical analysis was performed using ANOVA followed by Dunnett (**C**, and **E**) or Tukey (**J**) post hoc corrections, and unpaired two-tailed t test (**H**). \**p* < 0.05, \*\**p* < 0.01,\*\*\**p* < 0.001, and \*\*\*\**p* < 0.0001; ns, not significant.

### LAPTM4B inhibits the TGF-β/SMAD signaling pathway

The TGF-β/SMAD signaling pathway is crucial in mediating pro-fibrotic response across various cell types. To elucidate the molecular mechanism underlying the influence of LAPTM4B on the homeostasis of epithelial and fibroblast cells, we aim to investigate its effect on the TGF-β/SMAD signaling pathway. Upon shRNA-mediated knockdown of endogenous LAPTM4B, we observed a significant increase in TGF-β1-mediated rapid phosphorylation of SMAD2 in stable LAPTM4B-knockdown A549 cell lines (Figure 4A&B). This finding was further confirmed by siRNA-mediated LAPTM4B-knockdown in MRC-5 cells (Figure 4C&D). Conversely, transient overexpression of LAPTM4B in cells suppressed TGF-β1-induced phosphorylation of SMAD2 (Figure 4E&F). Additionally, AKT activation was not affected in LAPTM4B-knockdown or overexpressed A549 after TGF-β1 stimulation (Supplementary Fig. 3A&B). Immunofluorescence analysis provided additional evidence showing that shRNA-mediated knockdown of LAPTM4B expression resulted in spontaneous nuclear translocation of ectopically expressed HA-SMAD3 (Figure 4G). In contrast, ectopic expression of LAPTM4B suppressed TGF-β1-mediated nuclear translocation of SMAD3 in A549 cells (Figure 4H). Accordingly, siRNA-mediated knockdown of LAPTM4B increased TGF-β-inducible Smad binding element (SBE) luciferase activity (Figure 4I), whereas LAPTM4B overexpression suppressed TGF-β-induced SBE activity (Figure 4J).

**Figure 4.**
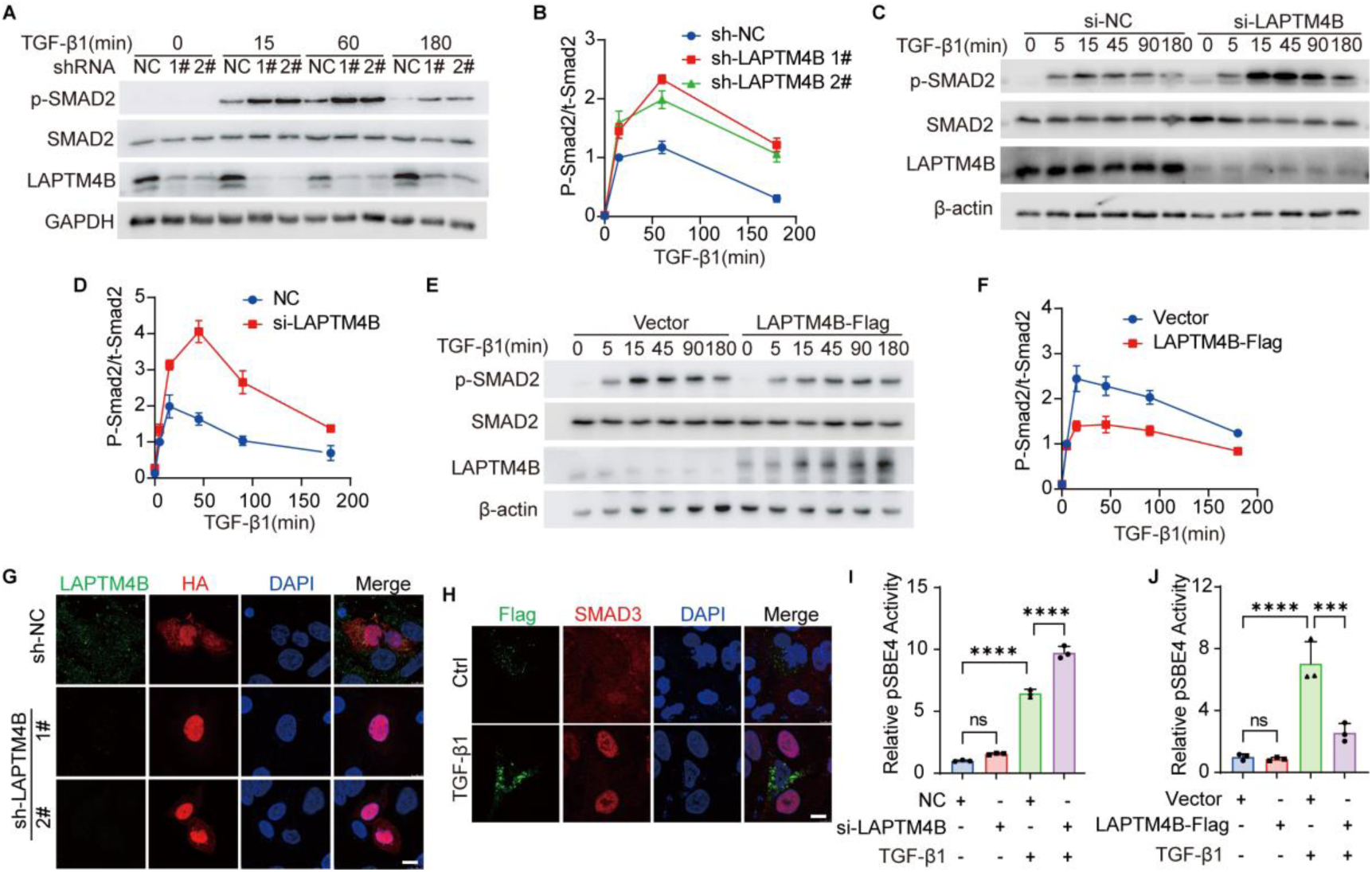
LAPTM4B dampens the TGFβ/SMADs signaling pathway. (**A**) Western blot analysis of p-SMAD2, total SMAD2, and LAPTM4B in sh-NC and sh-LAPTM4B A549 cells treated with TGF-β1 (5 ng/mL) for indicated durations. GAPDH served as loading control. (**B**) Relative phosphorylation levels of SMAD2, represented as p-SMAD2/total SMAD2 (mean ± S.D., n = 3). (**C**) Western blot analysis of p-SMAD2, total SMAD2, and LAPTM4B in MRC-5 cells transfected with si-NC or si-LAPTM4B for 48 h, followed by TGF-β1 (5 ng/mL) treatment for indicated durations. β-Actin served as loading control. (**D**) Relative phosphorylation levels of SMAD2, represented as p-SMAD2/total SMAD2, were quantified (mean ± S.D., n = 3). (**E**) Western blot analysis of p-SMAD2, total SMAD2, and LAPTM4B in MRC-5 cells transfected with LAPTM4B-Flag for 48 h, followed by TGF-β1 (5 ng/mL) treatment for indicated durations, β-actin served as loading control. (**F**) Relative phosphorylation levels of SMAD2, represented as p-SMAD2/total SMAD2 (mean ± S.D., n = 3). (**G**) Representative confocal images of immunofluorescence staining showing LAPTM4B (green) and HA-tag (red) in sh-NC or sh-LAPTM4B A549 cells transfected with HA-SMAD3 overexpression plasmid for 48 hours. DAPI was used to label cell nuclei, scale bar: 10 μm. (**H**) Representative confocal images of immunofluorescence staining showing Flag-tag (green) and SMAD3 (red) in A549 cells transfected with LAPTM4B-Flag for 48 hours followed by treatment with or without 5 ng/mL TGF-β1 for an additional 30 min. DAPI was used to label cell nuclei, scale bar: 10 μm. (**I**) Relative pSBE4 luciferase activity in MRC-5 cells transfected with either si-NC or si-LAPTM4B, followed by followed by treatment with or without 5 ng/mL TGF-β1 (mean ± S.D., n = 3). (**J**) Relative pSBE4 luciferase activity in MRC-5 cells transfected with either vector control or LAPTM4B-Flag, followed by treatment with or without 5 ng/mL TGF-β1 (mean ±S.D., n = 3). All statistical analysis was performed using ANOVA followed by Tukey post hoc corrections. \*\*\**p* < 0.001, and \*\*\*\**p* < 0.0001; ns, not significant.

### LAPTM4B interacts with TGFBR2 and destabilizes active SMAD2/3

Consistently with previous reports (26,34), we observed that Flag-tagged LAPTM4B partially overlaps with the early endosome marker EEA1 and co-localizes with the late endosome/lysosome marker RAB7 (Supplementary Fig. 4A&B). Given that the intracellular trafficking of the TGF-β receptor is known to influence downstream signaling dynamics, we aim to investigate whether LAPTM4B affects the intracellular trafficking of TGF-β receptors. Remarkably, TGFBR2-HA exhibited a high degree of co-localization with endogenous LAPTM4B that nearly all vesicles containing LAPTM4B also labeled for TGFBR2-HA (Figure. 5A). Interestingly, LAPTM4B expression in cells was stochastic, with some cells displaying high levels and others low levels of LAPTM4B. Notably, TGFBR2-HA predominantly localized to vesicles in cells with high LAPTM4B levels, rather than being distributed in the plasma membrane and cytoplasm as seen in cells with low LAPTM4B levels. Furthermore, TGF-β1 stimulation has no effect on the rates of TGFBR2-HA internalization into LAPTM4B-positive vesicles (Figure. 5A). The co-localization between LAPTM4B and TGFBR2 was further confirmed by transient co-transfection of LAPTM4B-Flag and TGFBR2-HA constructs and visualization using N-SIM microscopy and revealed that the co-localization of the two proteins occurred not only on the plasma membrane (Figure. 5B, inset 1) but also within LAPTM4B-containing vesicles (Figure. 5B, insets 1 and 2). Furthermore, immunoprecipitation assay confirmed the physical interaction between LAPTM4B and TGFBR2, irrespective of TGF-β1 stimulation (Figure 5C). In addition, SMAD3 was recruited to LAPTM4B-positive vesicles in the presence of TGF-β1 (Figure 5D, Supplementary Figure. S4C). Furthermore, LAPTM4B promoted the ubiquitination of SMAD3 under TGF-β1 stimulation (Figure 5E). The proteasome inhibitor MG132 restored the reduced phosphorylation of SMAD2 in response to LAPTM4B overexpression (Figure 5F&G). In addition, in vitro ubiquitination assays using ubiquitin mutants (K48O or K63O) indicated that LAPTM4B promoted K48-linked ubiquitination of both TGFBR2 (Figure 5H) and SMAD3 (Figure 5I) in the presence of TGF-β1.

**Figure 5.**
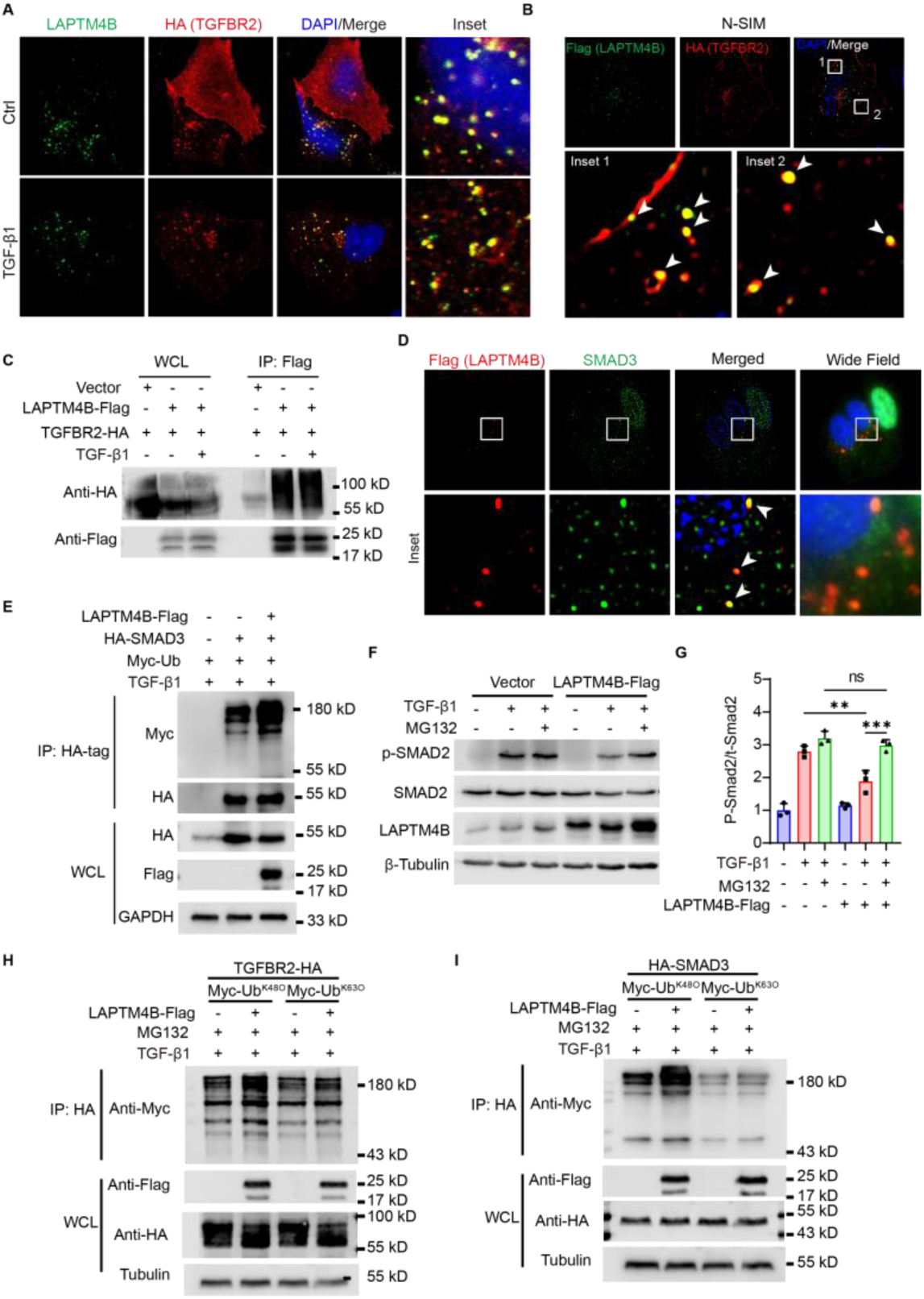
LAPTM4B interacts with and destabilizes TGFBR2 and SMAD3. (**A**) A549 cells were transfected with TGFBR2-HA for 48 hours, followed by treatment with or without 5 ng/mL TGF-β1 for an additional 30 min. Representative confocal microscopy images of LAPTM4B (green) and HA-tag (red), with cell nuclei stained with DAPI (blue). Detailed views of the boxed regions are shown. Scale bar: 5 μm. (**B**) A549 cells were co-transfected with LAPTM4B-Flag and TGFBR2-HA expression plasmids for 48 h. Representative N-SIM images of Flag (green) and HA-tag (red), with cell nuclei stained with DAPI (blue). Detailed views of the boxed regions are shown. Arrow heads indicate overlapped areas. (**C**) 293T cells were transfected with LAPTM4B-Flag and TGFBR2-HA, followed by treatment with or without 5 ng/mL TGF-β1 for an additional 30 min. Whole-cell lysates were immunoprecipitated with anti-Flag beads. (**D**) A549 cells were transfected with LAPTM4B-Flag for 48 hours, followed by treatment with 5 ng/mL TGF-β1 for an additional 30 min. Representative N-SIM images of Flag (red) and SMAD3 (green), with cell nuclei stained with DAPI (blue). Detailed views of the boxed regions are shown. (**E**) 293T cells were transfected with LAPTM4B-Flag, HA-SMAD3, and Myc-tagged ubiquitin, followed by treatment with 5 ng/mL TGF-β1 for an additional 30 min. Whole-cell lysates were immunoprecipitated with anti-HA beads. (**F**) Western blot and (**G**) densitometry analysis of p-SMAD2, total SMAD2, and LAPTM4B in A549 cells were transfected with LAPTM4B-Flag or vector for 48 hours, followed by treatment with MG132 (2 μM) for 2 hours before stimulation with 5 ng/mL TGF-β1 for an additional 30 min. β-Tubulin served as a loading control (mean ± S.D., n = 3). (**H**) 293T cells were transfected with LAPTM4B-Flag and TGFBR2-HA, or (**I**) HA-SMAD3, along with Myc-tagged Ub^K48O^ or Myc-tagged Ub^K63O^, and incubated for 48 h. Subsequently, the cells were treated with 5 ng/mL TGF-β1 for an additional 30 min, followed by whole-cell lysis and immunoprecipitation with anti-HA beads. Statistical analysis was performed using ANOVA followed by Tukey post hoc corrections (**G**). \*\**p* < 0.01, \*\*\*\**p* < 0.0001; ns, not significant.

### LAPTM4B-mediated recruitment of NEDD4L to endosomes facilitates ubiquitination of active SMAD2/3

The LAPTM family member LAPTM5 has previously been shown to interact with NEDD4 like E3 ubiquitin protein ligase (NEDD4L) through its C-terminal PY motifs(35). Since PY motifs are also present in LAPTM4B, we aim to investigate their potential to bind NEDD4L. Indeed, despite being predominantly distributed in the cytoplasm, HA-NEDD4L was also observed in LAPTM4B-Flag-positive puncta (Figure 6A). The co-localization between LAPTM4B and NEDD4L was further confirmed by visualizing exogenously expressed LAPTM4B-Flag and endogenous NEDD4L using N-SIM microscopy (Figure 6B). Co-immunoprecipitation (Co-IP) assay using HEK293T cells transiently co-transfected with HA-tagged NEDD4L and Flag-tagged LAPTM4B indicated that LAPTM4B physically binds to NEDD4L (Figure 6C). Furthermore, transfection with the HA-NEDD4L construct did not affect the levels of endogenous LAPTM4B, suggesting that LAPTM4B may not serve as a substrate for NEDD4L (Supplementary Figure S4A). Similar to LAPTM4B, overexpression of NEDD4L increased the ubiquitination level of SMAD3 under TGF-β1 stimulation (Figure 6D). Exogenous expression of NEDD4L decreased TGF-β1-induced SMAD2 phosphorylation in A549 cells (Figure 6E). Conversely, depletion of NEDD4L by siRNA transfection enhanced TGF-β1-induced SMAD2 phosphorylation (Figure 6F). Additionally, depletion of NEDD4L by siRNA transfection restored the suppressed phosphorylation level of SMAD2 (Figure 6G&H) and the activity of SBE luciferase reporter (Figure 6I) induced by LAPTM4B overexpression after TGF-β1 stimulation. Similarly, knockdown of LAPTM4B by siRNA restored the reduced phosphorylation level of SMAD2 (Figure 6J&K) and the activity of SBE (Figure 6L) induced by NEDD4L overexpression.

**Figure 6.**
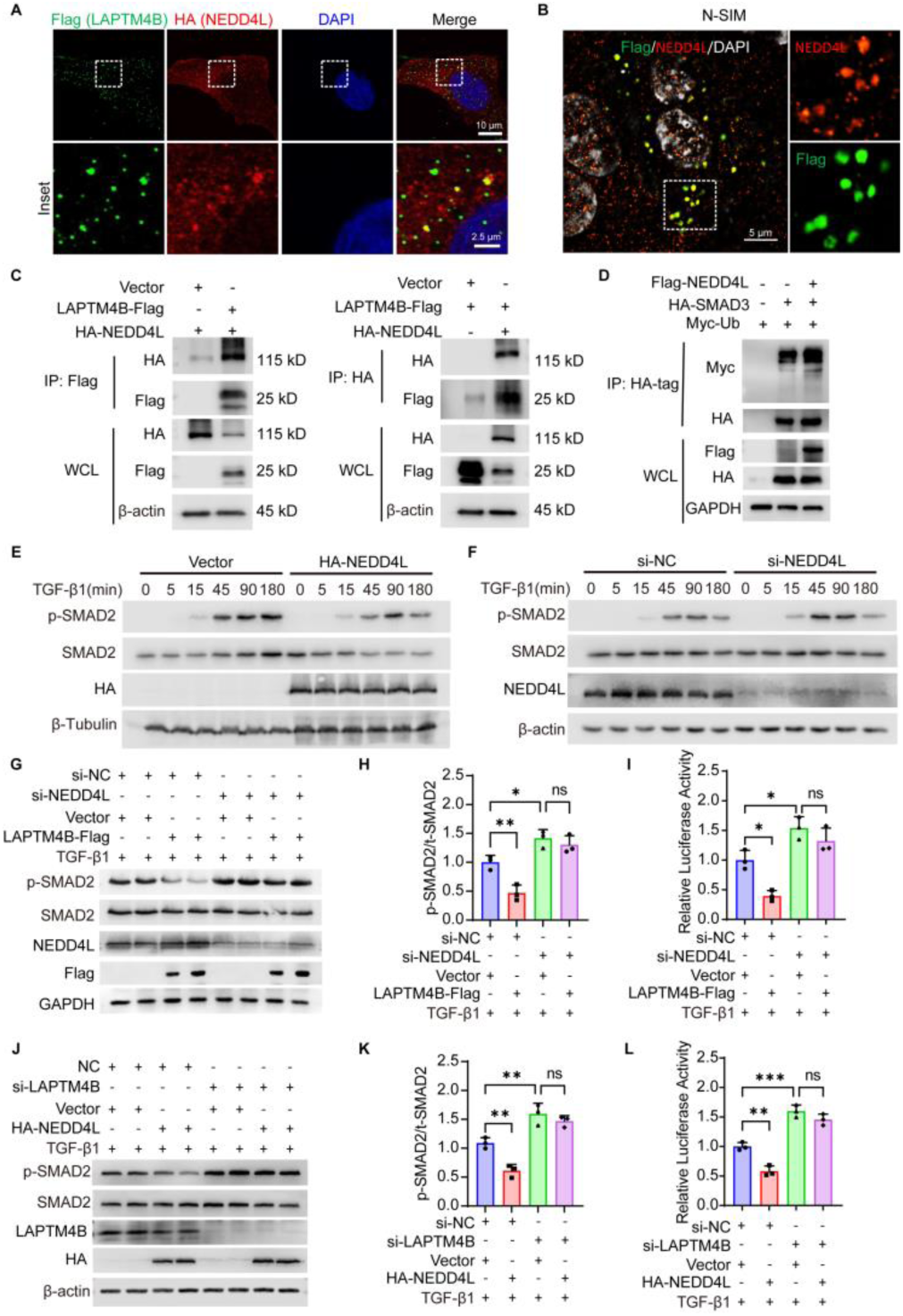
LAPTM4B-mediated recruitment of NEDD4L to endosomes facilitates ubiquitination of active TGFBR2 and SMAD3. (**A**) Confocal images of immunofluorescence staining for LAPTM4B-Flag (green) and HA-NEDD4L (red) in A549 cells, with cell nuclei stained with DAPI (blue). Scale bar: 10 μm. Detailed views of the boxed regions are shown. (**B**) N-SIM images of immunofluorescence staining for LAPTM4B-Flag (green) and endogenous NEDD4L (red) in A549 cells. Representative N-SIM images show Flag (green) and HA-tag (red), with cell nuclei stained with DAPI (gray). Detailed views of the boxed regions are shown. (**C**) 293T cells were transfected with LAPTM4B-Flag and HA-NEDD4L, and whole-cell lysates were immunoprecipitated with anti-Flag or anti-HA beads. (**D**) 293T cells were transfected with Flag-NEDD4L, HA-SMAD3, and Myc-tagged ubiquitin, followed by treatment with 5 ng/mL TGF-β1 for an additional 30 min. Whole-cell lysates were immunoprecipitated with anti-HA beads. (**E**) Western blot analysis of p-SMAD2, total SMAD2, and HA in A549 cells transfected with vector or HA-NEDD4L for 48 hours, followed by TGF-β1 (5 ng/mL) treatment for the indicated durations. β-Tubulin served as a loading control. (**F**) Western blot analysis of p-SMAD2, total SMAD2, and NEDD4L in A549 cells transfected with si-NC or si-NEDD4L for 48 hours, followed by TGF-β1 (5 ng/mL) treatment for the indicated durations. β-actin served as a loading control. (**G**) Western blot and (**H**) densitometry analysis of p-SMAD2, total SMAD2, NEDD4L, and Flag-tag in A549 cells transfected with the indicated siRNAs and plasmids for 48 hours, followed by TGF-β1 (5 ng/mL) treatment for 30 min. GAPDH served as a loading control (mean ± S.D., n = 3). (**I**) Relative pSBE4 luciferase activity in A549 cells transfected with the indicated siRNAs and plasmids, followed by TGF-β1 treatment (mean ± S.D., n = 3). (**J**) Western blot and (**K**) densitometry analysis of p-SMAD2, total SMAD2, LAPTM4B, and HA-tag in A549 cells transfected with the indicated siRNAs and plasmids for 48 hours, followed by TGF-β1 (5 ng/mL) treatment for 30 min. β-actin served as a loading control. (**I**) Relative pSBE4 luciferase activity in A549 cells transfected with the indicated siRNAs and plasmids, followed by TGF-β1 treatment (mean ± S.D., n = 3). All statistical analysis was performed using ANOVA followed by Tukey post hoc corrections. \**p* < 0.05, \*\**p* < 0.01, \*\*\**p* < 0.001; ns, not significant.

### AAV-mediated overexpression of Laptm4b alleviates BLM-induced lung fibrosis

Finally, the therapeutic potential of LAPTM4B was evaluated in the bleomycin-induced mouse model of lung fibrosis. Mice were administered adenovirus overexpressing Laptm4b (AAV-Laptm4b), while adenovirus expressing green fluorescent protein (AAV-Vector) served as the control. Fourteen days later, the mice were challenged with BLM for an additional 14 days (Figure 7A). Mice exposed to AAV-Laptm4b exhibited less weight loss following the BLM challenge (Figure 7B). Micro-CT images showed that AAV-Laptm4b-treated mice had a marked improvement in lung density after the BLM challenge (Figure 7C). Additionally, H&E and Masson’s trichrome staining demonstrated that mice treated with AAV-Laptm4b had better-preserved lung architecture compared to those treated with AAV-Vector (Figure 7D). These mice also showed significant reductions in Ashcroft scores (Figure 7E), inflammatory cells in BALF (Figure 7F), lung weight-to-body weight ratios (Figure 7G), and lung hydroxyproline levels (Figure 7H) after the BLM challenge. Western blot analysis confirmed the overexpression of LAPTM4B and the reduction of Fibronectin, Collagen1, and α-SMA expression in the lungs of AAV-Laptm4b-treated mice following the BLM challenge (Figure 7I&J). Collectively, these findings strongly suggest that LAPTM4B holds promise as a therapeutic target for the treatment of pulmonary fibrosis.

**Figure 7.**
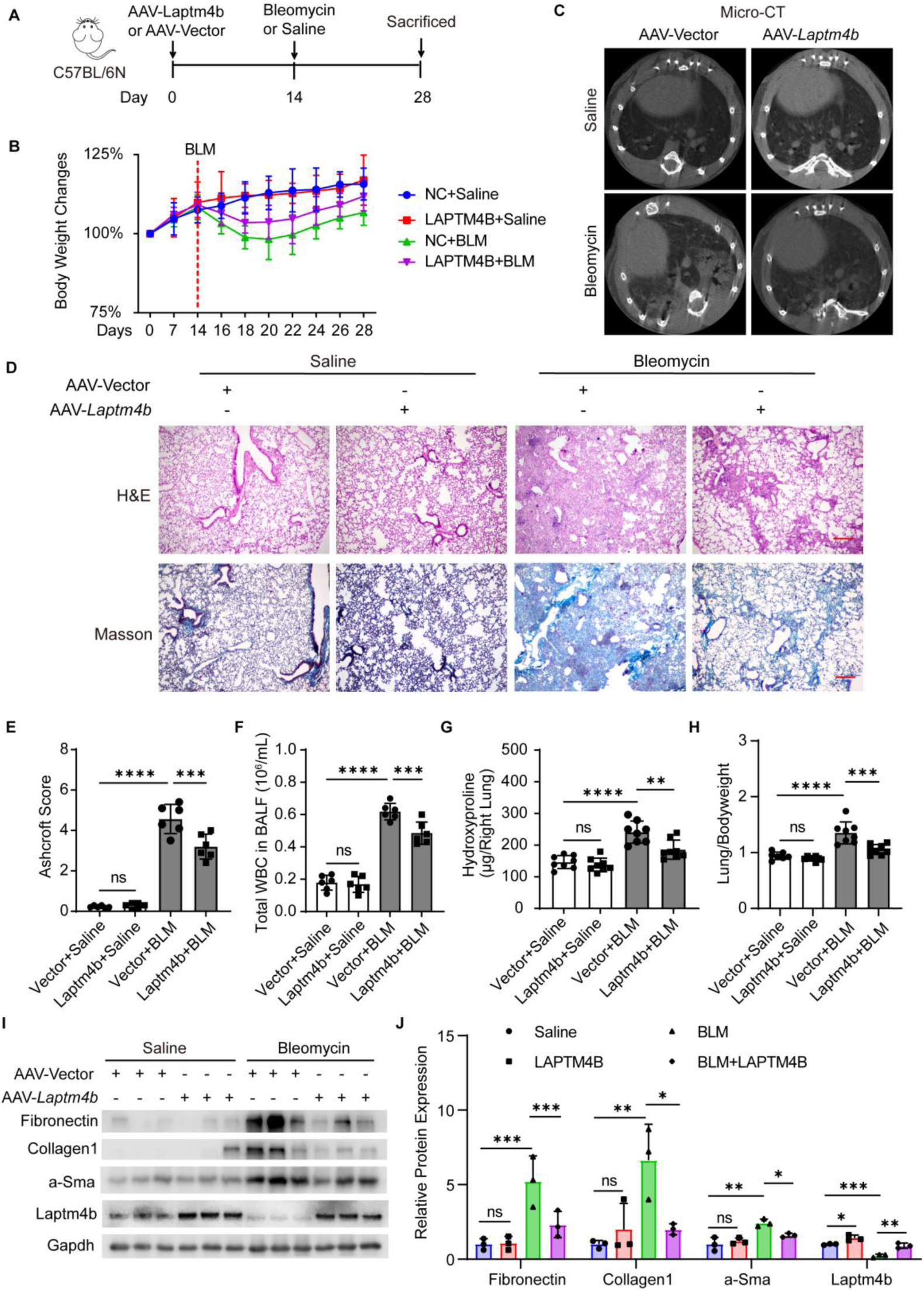
AAV-mediated LAPTM4B overexpression attenuates BLM-induced lung fibrosis. (**A**) Schematic diagram illustrating the experimental design. (**B**) Body weight changes of AAV Laptm4b or AAV-vector-infected mice treated with saline control or BLM (mean ±SD, n = 8 mice in Saline group, and 10 mice per group in other groups). (**C**) Representative Micro-CT images of mouse lungs (**D**) Hematoxylin and eosin (H&E) staining and Masson’s trichrome staining of lung sections (scale bar = 200 μm). (**E**) Ashcroft score analysis of H&E-stained lung sections (mean ±S.D, n = 6). (**F**) Quantification of the total number of WBC in BALF (mean ± SD, n = 6). (**G**) Mouse lung wet weight vs body weight (mean ± S.D., n = 8). (**H**) Measurement of hydroxyproline content in the right lungs (mean ±S.D., n = 8). (**I**) Western blot and (**J**) quantification of Fibronectin, Collagen1, α-Sma and Laptm4b levels in mouse lung homogenates (mean ±S.D., n = 3), Gapdh was used as loading control. All statistical analysis was performed using ANOVA followed by Tukey post hoc corrections. \**p* < 0.05, \*\**p* < 0.01,\*\*\**p* < 0.001, and \*\*\*\**p* < 0.0001; ns, not significant.

## Discussion

IPF is a chronic and progressive interstitial lung disease characterized by excessive deposition of extracellular matrix, leading to lung scarring and respiratory impairment. Despite extensive research, the underlying molecular mechanisms driving IPF pathogenesis remain incompletely understood. In this study, the role of LAPTM4B in modulating the TGF-β/SMAD signaling pathway in the context of IPF was investigated. Present findings reveal a novel interaction between LAPTM4B and NEDD4L, shedding light on the regulatory mechanisms governing TGF-β-mediated fibrotic responses in IPF.

In the present study, LAPTM4B was found to be drastically reduced in the lung tissue of IPF subjects and in mice with bleomycin-induced lung fibrosis. This reduction was also observed in acute lung injury models induced by bleomycin or LPS, suggesting that the decrease in LAPTM4B may occur early after the initial injury and inflammation phase, and persist during disease progression. However, the mRNA level of LAPTM4B remained unaffected in these mouse models. Additionally, most bulk gene expression analysis datasets based on microarray or RNA-seq of IPF subjects did not show significant changes or yielded contradictory results regarding LAPTM4B mRNA levels. The inconsistent protein and mRNA levels of LAPTM4B and other LAPTM family members have been well documented in other diseases or animal models. For instance, in a murine model of myocardial ischemia/reperfusion, the level of Laptm4b protein, but not mRNA, in the heart was significantly downregulated compared to the sham control (30). Similarly, in Neonatal Rat Cardiac Myocytes (NRCMs) during hypoxia/reoxygenation, while protein level reduced significantly, the mRNA level of LAPTM4B remained unchanged (30). In another case, despite a significant decline at the protein level, the mRNA levels of LAPTM5 were not affected in both mouse model of Non-alcoholic steatohepatitis and palmitic acid stimulated hepatocytes (35). These observations suggest that the loss of LAPTM4B in pulmonary fibrosis, as in other injury-related diseases, is limited to the protein level rather than the transcription level. This phenomenon may be related to the rapid turnover nature of LAPTM family proteins.

The present study demonstrated that normal levels of LAPTM4B are crucial for maintaining homeostasis in lung-derived epithelial and fibroblast cells, and suppress TGF-β1-induced fibrotic responses in both cell types. The TGF-β/SMAD signaling pathway is the most prominent regulator of pro-fibrotic responses. In canonical TGF-β/SMAD signaling, TGF-β1 binds to TGFBR2, leading to the recruitment and phosphorylation of TGFBR1, which in turn phosphorylates downstream effector proteins known as SMADs (36). Once phosphorylated, SMAD2 and SMAD3 form complexes with SMAD4, translocate into the nucleus, and regulate the transcription of target genes involved in fibrogenesis, including Collagen, Fibronectin, and α-SMA (36). Dysregulation of this pathway has been implicated in the pathogenesis of several fibrotic disorders, particularly pulmonary fibrosis (37). Aberrant activation of the TGF-β/SMAD signaling pathway promotes myofibroblast differentiation, excessive ECM deposition, and ultimately tissue scarring. Recent research has found that the glycolytic enzyme pyruvate kinase M2 (PKM2) enhances TGF-β/SMAD signaling and promotes profibrotic responses by disrupting the binding between SMAD7 and TGFBR1, reducing the ubiquitination and degradation of TGFBR1(38). Another research demonstrated that thioredoxin domain containing 5 (TXNDC5), an ER protein disulfide isomerase, is highly upregulated in IPF lungs and promotes fibrogenesis by enhancing TGF-β1 signaling through direct binding with and stabilization of TGFBR1 in lung fibroblasts (39). These findings highlight the complex regulatory mechanisms underlying the TGF-β/SMAD pathway and underscore the potential for targeting these interactions to develop new therapeutic strategies for pulmonary fibrosis.

Ubiquitination plays a critical role in regulating the turnover and activity of key components of TGF-β/SMAD signaling pathway (40). For instance, ubiquitination of TGF-β receptors (TGFBR1 and TGFBR2) can regulate their stability and endocytic trafficking, thereby influencing the duration and intensity of TGF-β signaling. Moreover, ubiquitination of SMAD proteins can regulate their transcriptional activity, stability, and subcellular localization, affecting downstream gene expression and cellular responses to TGF-β. E3 ubiquitin ligases, such as Smurf1, WWP1, NEDD4L and Arkadia, have been implicated in the ubiquitination and degradation of SMAD proteins, thereby acting as important regulators of TGF-β signaling (41,42). The dynamic interplay between ubiquitination and deubiquitination processes finely tunes the activity of the TGF-β/SMAD pathway, contributing to the downstream cellular processes including fibrogenesis (40,43). Further investigation into the specific ubiquitination events and their functional consequences within the TGF-β/SMAD signaling cascade will provide deeper insights into the molecular mechanisms underlying its physiological and pathological roles.

Indeed, LAPTM4B has been implicated in both receptor intracellular sorting and ubiquitin-dependent protein degradation (22). For instance, it impedes EGFR sorting by promoting the ubiquitination of Hrs (ESCRT-0 subunit), thus hindering Hrs from binding to ubiquitinate EGFR (26). Consequently, this process effectively curtails EGF-induced EGFR intracellular sorting and lysosomal degradation, resulting in amplified and prolonged EGFR signaling (26). Moreover, a very recent study demonstrated LAPTM4B suppress the phosphorylation and ubiquitination degradation of Yes-associated protein (YAP) in hepatocellular carcinoma, thereby preserving the stemness of the tumor cells (28). In the present observation, LAPTM4B interacts with TGFBR2, augmenting ubiquitination of TGFBR2 and active SMAD2/3, thus limited TGF-β/SMAD signaling. Given LAPTM4B is lack of inherent ubiquitin ligase activity, its participation in ubiquitin-dependent protein degradation hinges on its interaction with E3 ubiquitin ligases. Notably, several ubiquitin E3 ligases, such as SMURF1, SMURF2, NEDD4L, and WWP1, have been implicated in directly ubiquitination of TGF-β receptors and SMAD2/3, thereby contributing to the regulation of the TGF-β/SMAD signaling pathway (44). Members of the LAPTM family, including LAPTM5, LAPTM4A, and LAPTM4B, have been demonstrated to direct interact with NEDD4 E3 ubiquitin ligase family proteins via their C-terminal PY motifs (34,35,45–47). In current study, we demonstrated the co-localization and direct binding between LAPTM4B and NEDD4L, and clarifying that LAPTM4B is not a substrate for NEDD4L.

Previous researches have demonstrated declined NEDD4L expression in lung tissues from both IPF patients and bleomycin-induced mouse model of lung fibrosis (48,49). Conditional deletion of Nedd4l in lung epithelial cells causes progressive pulmonary fibrosis in both neonatal and adult mice (48,49). Recent investigations have shown that NEDD4L exerts anti-fibrotic effects by facilitating the ubiquitination and degradation of β-catenin (50) and TGFBR2 (51) in lung fibroblasts. Similarly, we confirmed that NEDD4L reduced TGF-β1-induced phosphorylation of SMAD2/3 by promoting ubiquitination of phosphorylation of SMAD2/3.

Previous studies have indicated that NEDD4L is primarily localized to the cell membrane and cytoplasm. The current observation indicates that in addition to the cell membrane and cytoplasm, NEDD4L also accumulates in LAPTM4B-positive vesicular structures. Given our observation that TGFBR2 also localizes to LAPTM4B-positive vesicles and that SMAD2/3 aggregates in LAPTM4B-positive vesicles following TGF-β1 treatment, we speculated that LAPTM4B anchors NEDD4L and TGFBR2 on the membrane of endocytic vesicles, thereby reducing the spatial distance between NEDD4L and activated SMAD2/3. This may facilitates NEDD4L to more efficiently ubiquitinate the activated TGF-β receptor-SMAD2/3 complex, thereby negatively regulating the TGF-β/SMAD signaling pathway. This hypothesis was further supported by LAPTM4B and NEDD4L are functionally interdependent.

Despite being an attractive target for IPF therapy, few candidates targeting the TGF-β/SMAD signaling pathway have even reached early-phase clinical trials (37). This may attribute to the ubiquitous distribution of TGF-β receptors in various cell types and the vital role of TGF-β/SMAD in pleiotropic functions, including cellular proliferation, differentiation, and normal repair processes (36). Therefore, finding more precise cellular and molecular targets to intervene is urgently needed. Despite NEDD4L also being found reduced in IPF and experimental fibrosis, we demonstrated that the protein level of LAPTM4B was nearly null at the very early stage after the initial lung injury. Loss of LAPTM4B may represent a rapid-response NEDD4L-mediated TGF-β/SMAD signaling switch which activates lung repair processes. On the other hand, preserved LAPTM4B levels in undamaged tissue and organs may block the action of paracrine and circulating TGF-β, thereby preventing the onset of fibrogenesis in unwanted areas. Thus, forced expression of LAPTM4B may offer the possibility to hijack the pathologically enhanced TGF-β/SMAD signaling in fibrotic areas to limit disease progression and to avoid the risk of severe side effects associated with direct inhibition. This hypothesis is supported by the observation that overexpression of Laptm4b exert therapeutic effects in bleomycin-induced mouse lung fibrosis.

This study has certain limitations that need to be addressed. Our in vivo research relied on the single-dose bleomycin mouse model, while this model closely mimics various pathological processes and characteristics observed in IPF, it cannot fully replicate the complexity of the disease due to our limited understanding of its mechanisms and inherent species differences between mice and humans (52). Moreover, although it has been demonstrated that tracheal instillation significantly reduces the systemic effects of AAV outside the lung, this cannot be completely ruled out (53). Additionally, although our study explored the interplay between LAPTM4B and NEDD4L, the specific mechanisms underlying this interaction, particularly the role of PY motifs, have not been independently verified.

In conclusion, we have elucidated a novel role of LAPTM4B in modulating the TGF-β/SMAD signaling pathway through its interaction with NEDD4L. This interaction impacts the ubiquitination and proteasomal degradation of key signaling molecules such as TGFBR2 and phosphorylated SMAD2/3, thereby regulating downstream responses associated with pulmonary fibrosis. These findings are of high interest because it may provide a possibility to curtail TGF-β-mediated fibrogenesis with minimum impact to healthy tissue. Further exploration of to identify the upper regulative mechanism of LAPTM4B level in repose to lung injury and fibrosis will add valuable insights into the complete LAPTM4B-NEDD4L-TGF-β/SMAD axis of IPF and may identify potential therapeutic targets IPF.

## Methods

### Reagents and antibodies. Recombinant human

TGF-β1 (240-B) was procured from R&D Systems (Minneapolis, MN, USA). SB431542 (HY-10431), Chloroquine (HY-17589A), MG132 (HY-13259) were obtained from MCE (Shanghai, China). Bleomycin hydrochloride were obtained from Hisun Pharm (Taizhou, China). Antibodies used in this study included anti-HA-tag (#3724), anti-DYKDDDDK-Tag (#8146), anti-Myc-tag (#2276), anti-COL1A1 (#72026), anti-α-SMA (#19245), anti-N-cadherin (#13116), anti-E-cadherin (#14472), anti-β-Tubulin (#2128), anti-phospho-Ser465/467 SMAD2 (#18338), anti-phospho-Ser423/425 SMAD3 (#9520), anti-SMAD2 (#5339), anti-SMAD3 (#9523), Alexa Fluor 488-conjugated anti-mouse IgG (#4408), and Alexa Fluor 594-conjugated anti-rabbit IgG (#8889) sourced from Cell Signaling Technology (Shanghai, China). Additionally, anti-NEDD4L (13690-1-AP), anti-Fibronectin (15613-1-AP), and anti-Vimentin (10366-1-AP) were acquired from ProteinTech (Wuhan, China). The Rabbit polyclonal antibody to LAPTM4B (DF12416), anti-phospho-Akt (Ser473) (AF0016), anti-Akt (AF6261), anti-GAPDH (AF7021), and anti-β-actin (AF7018) were obtained from Affinity Biosciences (Changzhou, China). anti-LAPTM4B (ab242376), anti-AGER (ab216329), Horseradish peroxidase (HRP)-conjugated Goat anti-rabbit (#ab205718) and HRP-conjugated anti-mouse IgG (#ab6789) were sourced from Abcam (Shanghai, China).

### Human lung tissue samples

Human lung tissue samples were collected following the guidelines outlined in the ATS/ERS/JRS/ALAT Clinical Practice Guidelines (54) at Henan Provincial Chest Hospital. In accordance with these guidelines, IPF lung tissues were obtained from patients undergoing open lung biopsy, while control non-IPF lung tissue samples were sourced from healthy lung tissue of age-matched donors without interstitial lung diseases who were undergoing surgical procedures for unrelated conditions.

### Mice

Male wild-type C57BL/6 mice (8 weeks old, weighing 18–22 g) were procured from Charles River Laboratory Animal Technology Co., Ltd. (Beijing, China). Male mice are exclusively used in the study because previous research has consistently shown that BLM-induced lung injury and fibrosis are more pronounced in male mice. This is a well-documented phenomenon that has led to the common practice of using only one gender in most studies. The mice were housed in a specific pathogen-free environment with controlled temperature and humidity, and a 12-hour light/dark cycle, and were provided *ad libitum* access to food and water. After acclimating to the environment for 1 week, the mice were subjected to gene knockdown or overexpression experiments in vivo. For gene knockdown, the mice were anesthetized with isoflurane and challenged with adeno-associated virus serotype 2/9 (AAV2/9) adenoviral particles (50 μL, 1.8×10^^12^ Vg/mL, *i.t.*) containing plasmid vector (pAAV-U6-sh-*Laptm4b*-CMV-EGFP-WPRE, Ref Seq: NM_033521, target sequence: 5’-CCCGTTCTTCTGTTACCAGAT-3’) or equivalent adenoviral particles loaded with a non-target control vector (pAAV-U6-shRNA(NC2)-CMV-EGFP-WPRE, sequence: 5’-TTCTCCGAACGTGTCACGTAA-3’), obtained from OBiO Technology Co., Ltd. (Shanghai, China). For gene overexpression, 50 μL (2.0×10^^12^ Vg/mL, *i.t.*) AAV2/9 adenoviral particles containing plasmid vector (pAAV-CMV-MCS-*Laptm4b-*3×Flag-EF1-GdGreen-WPRE, Ref Seq: NM_033521), or equivalent adenoviral particles loaded with pAAV-CMV-MCS-EF1-GdGreen-WPRE as control were administered. For inducing fibrosis, the mice were anesthetized with isoflurane and administered bleomycin (50 µL, 1.5 U/kg.bw, *i.t.*) or equivalent 0.9% saline as previously described (55). Finally, 14 days after the BLM challenge, the mice were euthanized with 20% Ethyl Carbamate (800 mg/kg, *i.p.*) (Sigma-Aldrich, Missouri, USA), and bronchoalveolar lavage (BAL) fluid was collected via tracheal insertion, and lung samples were collected. For inducing acute lung injury (ALI), the mice were anesthetized and challenged with bleomycin (1.5 U/kg.bw, *i.t.*) or lipopolysaccharide (LPS) (4 mL/kg.bw, *i.t.*; Sigma-Aldrich), lung samples were collected 24 hours after the challenge. For marking ACE2s in vivo, *Sftpc*-CreERT2 and *Rosa26R-CAG-lsl*-tdTomato (*R26R*-tdTomato) mice were given 3-4 doses of tamoxifen (100 mg/kg, *i.p.*; Sigma-Aldrich).

### Micro-CT Imaging

Micro-computed tomography (Micro-CT) imaging was performed prior to euthanasia of the mice, following a previously described protocol (56). Briefly, the mice were anesthetized with isoflurane and positioned in the supine orientation. Micro-CT imaging was conducted using a SkyScan 1276 micro-CT system (Bruker, Kontich, Belgium) with scanning parameters set at 60 kV X-ray tube voltage, 200 µA anode current, and a 0.5 mm aluminum filter. Subsequently, the obtained images underwent reconstruction and superimposition using Insta-Recon software (Version 2.1.0.00).

### Lung histological analysis and immunohistochemical staining

For histological analysis, human lung tissue samples or mouse left lung lobes were fixed for 24 hours in 4% phosphate-buffered formalin (PFA), dehydrated using a gradient ethanol solution and xylene, and embedded in paraffin. The tissues were then sliced into 4 μm sections. Tissue slides were dewaxed with xylene, rehydrated with descending concentrations of ethanol solution, and rinsed with deionized water. Masson’s trichrome stain and H&E stain were performed using a Masson’s trichrome staining kit (G1340, Solarbio, Beijing, China) or an H&E staining kit (G1120, Solarbio), respectively, following the manufacturer’s protocols. For Ashcroft score analysis (57), five fields were randomly selected from each H&E-stained lung section under 100× magnification. The average score for each field was independently determined by two pathologists in a blinded manner. For immunohistochemical staining, endogenous peroxidase activity was blocked by incubating sections in a 3% hydrogen peroxide solution at room temperature for 30 min after antigen retrieval. The slides were then blocked with 10% normal goat serum for 1 hour. Primary antibodies were applied as indicated and incubated overnight at 4 °C, followed by incubation with an HRP-conjugated secondary antibody for 60 min at room temperature. Visualization was achieved by DAB staining, and counterstaining was performed with hematoxylin. Images were captured using an Olympus BX43 upright microscope (Tokyo, Japan).

### Hydroxyproline Assay

The quantification of hydroxyproline content in mouse lungs was conducted using a Hydroxyproline Assay Kit (MAK008, Sigma, St. Louis, MO, USA) following the manufacturer’s instructions. Initially, fresh right mouse lungs were homogenized in 100 μL of deionized water for every 10 mg of tissue. Subsequently, an equal volume of 12 N HCl aqueous was added to the homogenate, and hydrolysis took place at 120°C for 3 hours in Teflon-capped vials. Following hydrolysis, the samples underwent centrifugation at 10,000 × *g* for 15 min to eliminate any residues. Next, 10 μL of each sample was carefully transferred to a 96-well plate and evaporated at 60°C until completely dry. Subsequently, 100 μL of the Chloramine T/Oxidation buffer mixture was added to each sample, followed by incubation at room temperature for 5 min. This was followed by the addition of p-Dimethylaminobenzaldehyde reagent (100 μL), and the mixture was further incubated for 90 min at 60°C. The absorbance at λ=560 nm of each well was measured using a BioTek ELX800 plate reader (BioTek, Winooski, VT, USA). The hydroxyproline content in the samples was calculated based on the standard curve plotted using the hydroxyproline standard. The data were presented as µg of hydroxyproline per right lung.

### Cell culture, and transfection

Human non-small cell lung cancer cell line A549 was obtained from Procell (Wuhan, China) and cultured in DMEM/F12 medium (PM150310, Procell, Wuhan China), supplemented with 10% FBS (Gibco, Grand Island, NY), 100 U/mL penicillin, and 100 μg/mL streptomycin (Solarbio, Beijing, China). The human embryonic lung fibroblast cell line MRC-5, obtained from ATCC, was cultured in MEM (Hyclone) supplemented with 10% FBS, 2 mM L-glutamine, 100 U/mL penicillin, and 100 μg/mL streptomycin (Solarbio, Beijing, China). HEK293T cells, also obtained from Procell (Wuhan, China), were cultured in high-glucose DMEM (Hyclone) supplemented with 10% FBS, 2 mM L-glutamine, 100 U/mL penicillin, and 100 μg/mL streptomycin. All experiments involving these cell lines were conducted within 10 passages, and all cell lines tested negative for mycoplasma contamination. Cell cultures were maintained in an incubator (Thermo Fisher, Waltham, MA, USA) at 37°C with 5% CO_2_ and saturated humidity. The following plasmids were used: empty pcDNA3.1(+) plasmid (Vector), pcDNA3.1(+) expression plasmids encoding C-terminal Flag-tagged LAPTM4B, N-terminal HA-tagged NEDD4L, N-terminal Flag-tagged NEDD4L, N-terminal HA-tagged TGFBR2, and N-terminal HA-tagged SMAD3. Additionally, Myc-tagged wildtype ubiquitin and the K48O and K63O ubiquitin mutants were utilized. These plasmids were constructed using previously described methods(58). Plasmid or siRNA transfections in A549 and MRC-5 cells were performed using Lipofectamine 3000 (Invitrogen) or the riboFECT CP Transfection Kit (RiboBio, Guangzhou, China). Plasmid transfections in HEK293T cells were performed using Polyethylenimine Linear (PEI; 40816ES03, Yeasen Biotechnology Co., Ltd., Shanghai, China).

### Stable Cell Lines

To establish stable knockdown of LAPTM4B in A549 cells, we utilized independent small hairpin RNA (shRNA) sequences specifically designed to target LAPTM4B (Ref Seq: NM_033521). The sh-*LAPTM4B* 1# sequence (TRCN0000150679) had the target sequence 5’-GATGATGTCATGTCAGTGAAT-3’, while the sh-*LAPTM4B* 2# sequence (TRCN0000150864) had the target sequence 5’-GATATGTGCTATGGCTACTTA-3’. These shRNA sequences were incorporated into the pLKO.1 vector. As a control, the pLKO.1 vector expressing a non-targeting shRNA (SHC016, Sigma-Aldrich) was used (sh-NC). The transfection of lentiviral plasmids containing the shRNA sequences, along with packaging plasmids (psPAX2 and pMD2.G), was carried out using PEI on 293T cells. After 48-72 hours post-transfection, the lentiviral supernatant containing viral particles was collected and subsequently filtered through a 0.45 µm filter to remove cell debris. A549 cells were transduced with shRNA lentiviral particles in the presence of 5 mg/mL polybrene. Following transduction, positive selection was performed using puromycin (2 μg/mL; HY-B1743A, MCE) for duration of 7 days. Subsequently, the cells were maintained in completed medium supplemented with 2 μg/mL puromycin. This rigorous selection and maintenance process ensured the establishment of stable cell lines exhibiting LAPTM4B knockdown for further experimentation.

### Immunofluorescence staining

Cells were seeded on poly-L-lysine-coated glass coverslips in a 24-wells plate. After indicated treatments, cells were fixed with 4% paraformaldehyde in PBS for 30 min at room temperature. Cells were then washed with PBS and subsequently permeabilized with 0.2% Triton-100 in PBS for 10 min and then blocked with 5% goat serum and 1% BSA in PBS for 1 h at room temperature. The coverslips were then incubated overnight at 4 °C with primary antibodies followed by incubated for 1 hour at room temperature with Alexa Fluor 488- and/or Alexa Fluor 568-, and/or Alexa Fluor 647-conjugated secondary antibodies, and cell nuclei were stained with DAPI (C0065, Solarbio, Beijing, China). Subsequently, the coverslips were mounted on glass slides with anti-fading mounting medium (S2100, Solarbio, Beijing, China). Images were obtained using a laser scanning confocal microscope (Leica, Wetzlar, Germany) or a Super-Resolution Microscope System (Nikon Instruments Inc. Melville, NY, USA.).

### Western blot analysis

The Western blot analyses were conducted following established procedures (53). Briefly, cell lysates or lung homogenates were prepared using RIPA lysis buffer (P0013B, Beyotime, Shanghai, China), supplemented with protease and phosphatase inhibitors. Following sonication and centrifugation at 10,000 × *g* for 15 min, the supernatants were collected and boiled at 95 °C for 5 min. Protein concentrations in the supernatants were determined using the BCA kit (PC0020, Solarbio, Beijing, China). Equal amounts of protein were loaded onto 8–12% SDS-PAGE gels and subjected to electrophoresis, followed by electrotransferring to PVDF membranes (Millipore, Burlington, MA, USA). The membranes were then blocked with 5% skim milk and incubated overnight at 4 ℃ with the indicated primary antibodies. Subsequently, the membranes were incubated with HRP-conjugated secondary antibodies for 1 hour at room temperature. Bands were visualized using enhanced chemiluminescence (Thermo Fisher, Waltham, MA, USA), and images were captured using an Odyssey Imaging System (LI-COR Biosciences, Lincoln, NE, USA). ImageJ version 1.54g (National Institutes of Health, Bethesda, MD, USA; https://imagej.net/ij/) was used for gray scale analysis of Western blot bands.

### Quantitative RT-PCR analysis

Cells were lysed using TRIzol (Takara, Dalian, China) to extract total RNA. cDNA was synthesized using the GoScript Reverse Transcription System (A5001, Promega, Madison, WI, USA). The reaction was performed by using using SYBR green kit (11201ES03, Yeasen) on a LightCycler 480 system (Roche, Penzberg, Germany). The primers used were as follows: Laptm4b forward, 5′-CCTGGACTCGGTTCTACTCCC-3′; reverse, 5′-CCCCTCCTAGTTCAGAGCCT-3′. Actb forward, 5′-CATTGCTGACAGGATGCAGAAGG-3′; reverse, 5′-CCACAGGATTCCATACCCAAG-3′. Relative mRNA levels were quantified by calculating the comparative 2^-ΔΔCt^ method.

### Collagen gel contraction assay

Briefly, MRC-5 cells were transfected with the indicated siRNA or plasmid for 24 hours before being suspended in MEM and mixed with rat tail collagen type 1 solution (354236, Corning, NY, USA) at a 2:1 ratio. The mixture was neutralized and added to a 24-well plate (0.5 mL/well), allowing it to polymerize at 37°C in a CO₂ incubator for 1 hour. After polymerization, 0.5 mL of culture medium with or without TGF-β1 (5 ng/mL) was added and incubated overnight. The gels were then released from the wells using a sterile scalpel. Images were taken after 24 hours, and were analyzed using ImageJ software (version 1.54g). The gel contraction rate was calculated as the contracted area divided by the initial gel surface area.

### Luciferase assays

Luciferase assays were performed as previously described(59). In brief, cells were transfected with the indicated plasmids or siRNAs, and pSBE4-TA-Luc reporter plasmid (D4320, Beyotime) for 24 hours and then serum starved overnight before treatment with TGF-β1 (5 ng/mL). Cells were harvested 12 hours after stimulation to perform luciferase assays using a luciferase assay system (E1500, Promega) according to the instructions.

### Immunoprecipitation assays

Immunoprecipitation was performed to detect the interactions between proteins. Briefly, HEK293T cells were co-transfected with the indicated plasmids using PEI. After transfection for approximately 48 hours, cell was treated as indicate before were lysed using ice-cold WB/IP lysis buffer (20 mM Tris-HCl, pH 7.4; 150 mM NaCl; 1 mM EDTA; and 1% NP-40) containing a protease inhibitor cocktail (P1005; Beyotime, Shanghai, China). After centrifugation at 12,000 × *g* for 10 min, the supernatant was collected. Each sample was subsequently incubated with Anti-HA or Anti-Flag magnetic beads (P2121 or P2115, Beyotime) at 4 °C for 4 hours. After centrifugation (3,500 × *g* for 5 min), the beads were washed with PBS twice. Finally, the beads were heated at 95°C in SDS loading buffer for 10 min, and separated by SDS-PAGE for western blotting.

### Ubiquitination assays

HEK293T cells were seeded in 10 cm culture dishes and co-transfected with indicated plasmids for 48 hours. Subsequently, the cells were lysed in 600 μL WB/IP lysis buffer supplemented with 60 μL of 10% SDS lysis buffer containing a protease inhibitor cocktail, followed by sonication and denatured by heating at 95 °C for 10 min. After centrifugation at 10,000 × *g* for 10 min, the supernatant was collected and incubated with Anti-HA or Anti-Flag magnetic beads overnight at 4 °C. The beads were then removed after centrifugation at 3,500 × *g* for 5 min and washed three times with PBS buffer. Subsequently, the beads were heated at 95 °C with loading buffer for 10 min and separated by SDS-PAGE for western blotting, as previously described.

### Wound-healing assay

Cells were treated as indicated and subsequently transferred to a 24-well plate for continuous cultivation. Upon reaching 90–100% confluence, a linear scratch was generated using a sterile pipette tip. After removing cellular debris by washing with PBS, the scratch was documented at 0-hour and 24-hour intervals using an inverted microscope (Leica, Wetzlar, Germany). The ImageJ software was utilized to measure the wound width at each time point. The wound healing percentage was calculated using the formula:

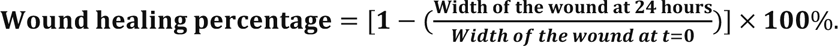

### Statistics

Normality tests were conducted for all data before analysis to determine whether parametric or nonparametric statistical tests were appropriate. The two-tailed Student’s t-test or Mann-Whitney U test was employed to compare two experimental groups, depending on the distribution of the data. For comparisons among multiple groups, one-way ANOVA followed by Tukey post hoc corrections was performed.

### Study approval

Prior to human lung sample collection, the research protocol was approved by the Henan Provincial Chest Hospital Medical Research Ethics Committee (No. 2019-05-07), and written informed consent was received from all patients undergoing surgical procedures prior to participation. The study strictly adhered to the principles outlined in The Code of Ethics of the World Medical Association (Declaration of Helsinki) governing experiments involving human subjects. Animal care and handling procedures were conducted in accordance with institutional and national guidelines and were approved by the Henan Normal University Institutional Animal Care and Use Committee.

## Data availability

The datasets used and/or analysed during the current study available from the corresponding author on reasonable request.

## Acknowledgements

This work was supported by the Ministry of Science and Technology, PR China grant 2019YFE0119500 (G.Y.), and Key R&D Program of Henan province grant 231111310400 (G.Y.); Henan Project of Science and Technology grants 232102521025 (L.W.), and GZS2023008 (L.W.).

