## Supplementary Figures for "LAPTM4B Alleviates Pulmonary Fibrosis by Enhancing NEDD4L-Mediated TGF-β Signaling Suppression"

Supplementary Fig1

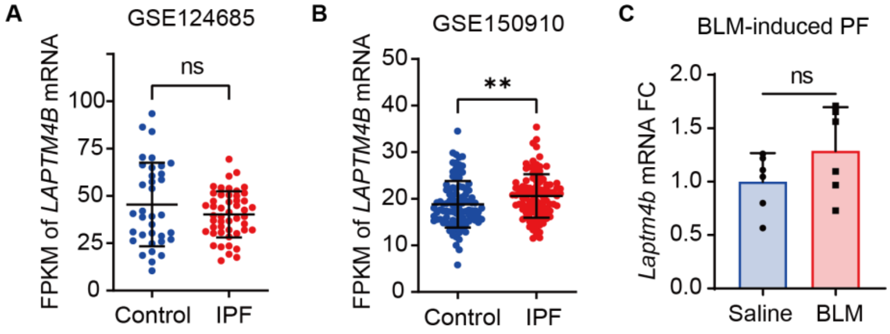


(**A**) Scatterplot showing the Fragments Per Kilobase of transcript per Million mapped reads (FPKM) of LAPTM4B mRNA in the GSE124685 RNA-seq dataset. Statistical analysis was performed using an unpaired two-tailed t-test (mean ± S.D., Control: n = 35, IPF: n = 49). (**B**) Scatterplot of the FPKM of LAPTM4B mRNA in the GSE150910 RNA-seq dataset. Statistical analysis was performed using an unpaired two-tailed Mann-Whitney test. (mean ± S.D., n = 103 subjects per group). (**C**) Quantitative RT-PCR analysis of Laptm4b mRNA levels in homogenates from bleomycin-induced fibrotic lungs. Acta1 was used as an internal control. Statistical analysis was performed using an unpaired two-tailed t-test (mean ± S.D., n = 6 mice per group). **$p < 0.01$; ns, not significant.

Supplementary Fig2

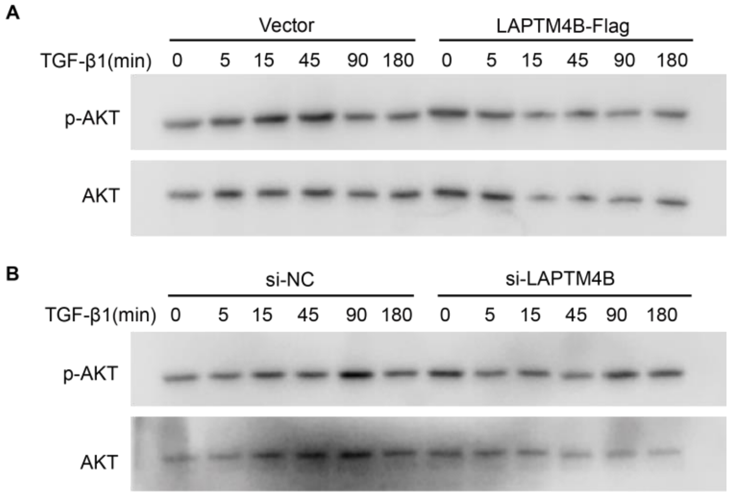


(**A**) Western blot analysis of p-AKT, and total in MRC-5 cells transfected with LAPTM4B-Flag for 48 h, followed by TGF-β1 (5 ng/mL) treatment for indicated durations. (**B**) Western blot analysis of p-AKT, and total AKT in MRC-5 cells transfected with si-NC or si-LAPTM4B for 48 h, followed by TGF-β1 (5 ng/mL) treatment for indicated durations.

Supplementary Fig3

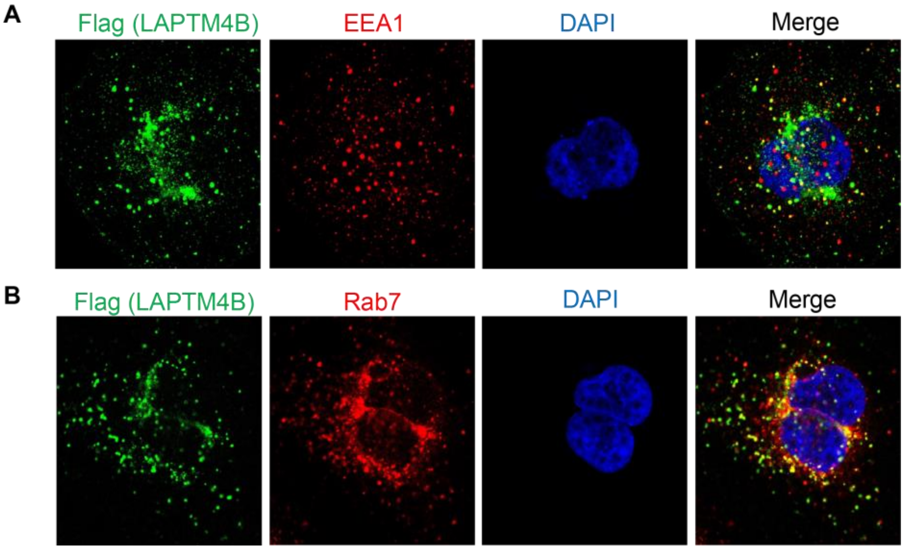


(A) A549 cells were transfected with LAPTM4B-Flag for 48 hours. Representative confocal microscopy images show LAPTM4B (green) and EEA1 (red), with cell nuclei stained with DAPI (blue). (B) A549 cells were transfected with LAPTM4B-Flag for 48 hours. Representative confocal microscopy images show LAPTM4B (green) and Rab7 (red), with cell nuclei stained with DAPI (blue).

Supplementary Fig4

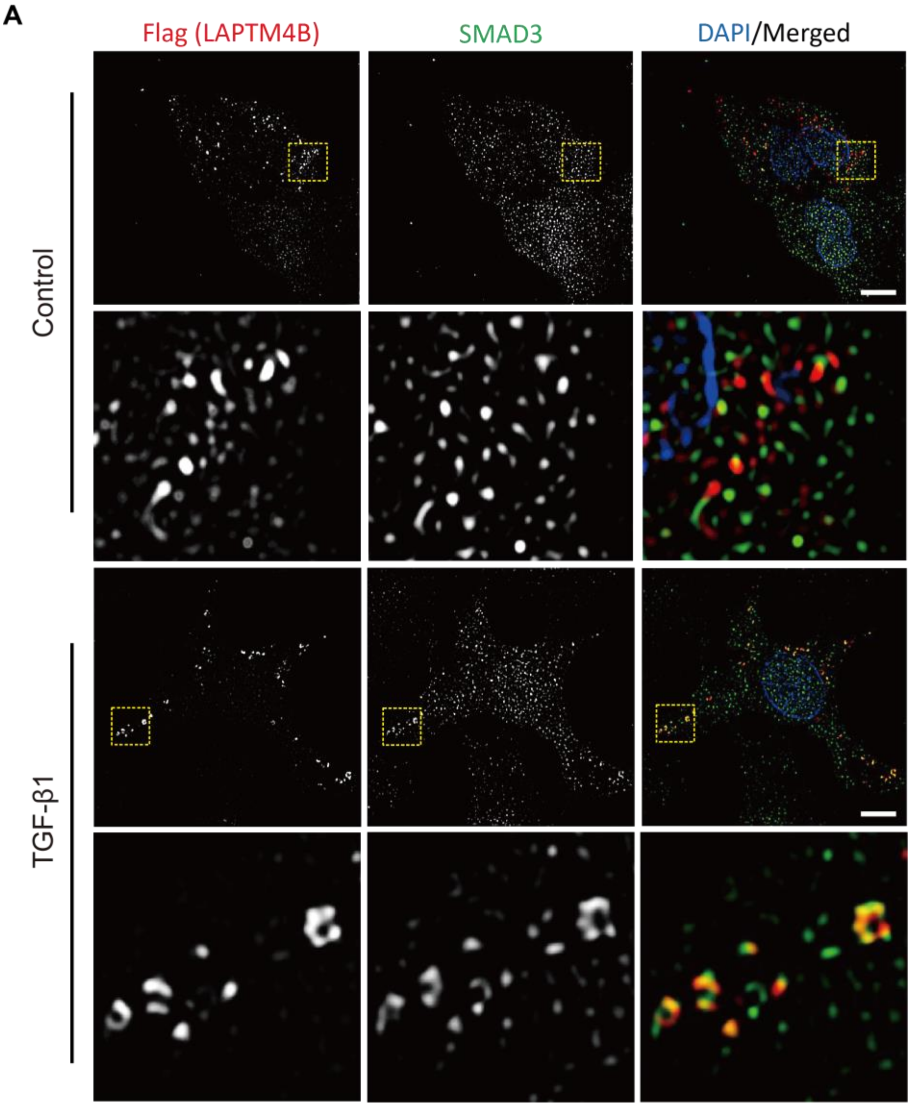


(**A**) A549 cells were transfected with LAPTM4B-Flag for 48 hours, followed by treatment with or without 5 ng/mL TGF-β1 for an additional 30 min. Representative N-SIM images of Flag (red) and SMAD3 (green), with cell nuclei stained with DAPI (blue).

Supplementary Fig5

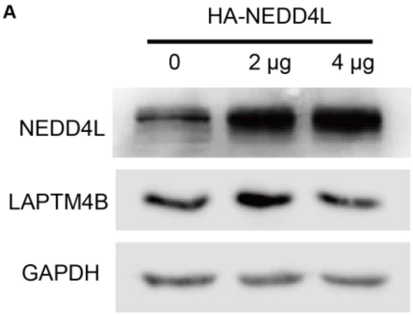


(**A**) Western blot analysis of NEDD4L, and LAPTM4B in A549 cells transfected with HA-NEDD4L for 48 h. GAPDH used as a loading control.
